# *Ex vivo* live cell tracking in kidney organoids using light sheet fluorescence microscopy

**DOI:** 10.1101/233114

**Authors:** Marie Held, Ilaria Santeramo, Bettina Wilm, Patricia Murray, Raphaël Lévy

## Abstract

Screening cells for their differentiation potential requires a combination of tissue culture models and imaging methods that allow for long-term tracking of the location and function of cells. Embryonic kidney re-aggregation *in vitro* assays have been established which allow for the monitoring of organotypic cell behaviour in re-aggregated and chimeric renal organoids. However, evaluation of cell integration is hampered by the high photonic load of standard fluorescence microscopy which poses challenges for imaging three-dimensional systems in real-time over a time course. Therefore, we employed light sheet microscopy, a technique that vastly reduces photobleaching and phototoxic effects. We have also developed a new method for culturing the re-aggregates which involves immersed culture, generating organoids which more closely reflect development *in vivo*. To facilitate imaging from various angles, we embedded the organoids in a freely rotatable hydrogel cylinder. Endpoint fixing and staining were performed to provide additional biomolecular information. We succeeded in imaging labelled cells within re-aggregated kidney organoids over 15 hours and tracking their fate while simultaneously monitoring the development of organotypic morphological structures. Our results show that Wt1-expressing embryonic kidney cells obtained from transgenic mice could integrate into re-aggregated chimeric kidney organoids and contribute to developing nephrons. Furthermore, the nascent proximal tubules that formed in the re-aggregated tissues using the new culture method displayed secretory function, as evidenced by their ability to secrete an organic anion mimic into the tubular lumen.

## Introduction

Culture systems for growing intact kidney rudiments *ex vivo*, first established by Grobstein in the 1950s, are excellent tools for understanding various aspects of kidney development (1, 2). In 2002, Atala’s group showed that dissociated bovine kidney rudiments had the remarkable ability to re-aggregate and generate nephron-like structures following subcutaneous implantation into adult cows (3). More recently, it has been shown that disssociated mouse kidney rudiments can re-aggreate and generate functional nephrons and ureteric buds when cultured *ex vivo* (4-6). These re-aggreated rudiments are valuable tools for determining the nephrogenic potential of different types of stem or progenitor cells, because the stem/progenitor cell type under investigation can be labelled and then mixed with the dissociated embryonic kidney cells to generate a chimeric re-aggregate (6-10). The recombinant kidney organ culture involves the isolation of the intact, developing organ, its dissociation and re-aggregation of the cell mixture, which is then typically cultured at the air-liquid interface atop a micro-porous filter.

In the developing kidney, reciprocal inductive interactions between the ureteric bud and metanephric mesenchyme lead to the formation of the metanephric kidney (11, 12). One of the key transient nephrogenic structures is the cap mesenchyme which, as a morphologically distinguishable layer 4-5 cells deep, is essential for inducing ureteric bud branching as well as providing a source of progenitor cells, ultimately giving rise to the whole nephron, including glomeruli and renal tubules (13, 14). Any system mimicking renal development should both closely resemble *in vivo* organogenesis, including the formation of important nephrogenic structures, and allow for close monitoring of events. Unfortunately, the re-aggregation assay, though reproducible, results in the generation of organoids that differ in size and organisation. This makes it difficult to generate and compare quantitative data between experiments. Longitudinal monitoring of a single experiment would therefore benefit our understanding of cellular and molecular events during kidney development. Furthermore, cell tracking can help us to answer previously unanswered questions. For instance, what proportion of the labelled cells survives, and when does cell death occur? Do the surviving cells integrate into developing structures? If so, do they actively migrate to developing structures? Previous studies have shown that various types of stem and progenitor cells can integrate into developing structures within chimeric re-aggregates *ex vivo* and can also exhibit function, but it is not known how and when they reach the respective renal structures (6, 7, 9, 15).

Previously, ureteric bud tip cells have been tracked manually in single focal plane confocal images (16). Lindström et al. (17) performed time-lapse capture of developing nephrons of a signalling reporter mouse strain but quantitative analysis was undertaken following fixation and immunofluorescent staining. Finally, Saarela at al. (18) recorded z-stacks of mouse embryonic kidney organoids repeatedly for up to 20 mins. Automated cell tracking of ureteric bud cells was however performed on single focal plane images.

Re-aggregated tissues cultured at the air-liquid interface grow/self-assemble primarily in two dimensions with little increase in depth (5). While a flat tissue is advantageous for standard optical microscopy imaging of simulated early kidney development, it does not accurately reflect kidney development *in vivo*, as the developing kidneys *in situ* are spheroidal structures. An organoid context would mimic the physiological situation better but poses greater imaging difficulties. For end point imaging, organoids/organ tissue spheroids are often cultured, fixed, sectioned, stained, and imaged (19). In such conditions, the three-dimensional morphological and general context are maintained during culture but lost for imaging.

Light sheet fluorescence microscopy (20) allows for the optical sectioning of the sample while maintaining its three-dimensional architecture, i.e. whole mount analysis (20-23). The imaging system (Zeiss Z.1 Lightsheet) combines two-dimensional illumination with orthogonal camera-based detection. A cylindrical lens shapes the laser light into a thin sheet of light directed onto the sample, illuminating only a section. The camera-based detector records the whole focal plane at once, resulting in fast imaging times. With only a section of the sample being illuminated at any time and rapid frame-wise data capture, light sheet fluorescence microscopy creates a photonic load several orders of magnitude lower than standard confocal fluorescence imaging (20), therefore allowing the capture of transient phenomena (16-18, 24, 25). For Samples are suspended in a water-based gel and can be rotated freely around one axis and therefore imaged from various angles. For live imaging, the samples in the hydrogel are surrounded by growth medium and a controlled environment (temperature, CO2 concentration). Cell spheroids have repeatedly been imaged using light sheet fluorescence microscopy (23, 25-28). Furthermore, the imaging technique has been used to investigate dynamic processes on varying scales, including tracking microtubules plus tips of the mitotic apparatus (29), lineage tracing of cells in spheroids (24) up to *in toto* imaging of mouse cells from zygote to blastocyst (30) as well as whole embryos of *Drosophila melanogaster* (31) and zebrafish (32), further advancing knowledge about developmental processes. These works all involved automated tracking algorithms.

Here, we have adapted the re-aggregation assay, creating embryonic renal organoids rather than flat tissues. This is the first time that re-aggregated kidney rudiments have been imaged by light sheet fluorescence microscopy. Performing long-term time lapse imaging, we aimed to track labelled cells during the culture period, and subsequently analyse the tracking data.

## Materials and Methods

### Materials

Materials were purchased from Sigma-Aldrich (Sigma-Aldrich Corp., USA) unless otherwise indicated.

### PDMS-polycarbonate membrane well fabrication

A custom organoid well construct was fabricated using the transparent, biocompatible polymer PDMS (Sylgard 184, Dow Corning, USA) (33). Briefly, the PDMS pre-polymer mixed with its curing agent (10:1 by weight) was degassed, poured into 35mm Petri dishes and cured at 60°C on a hotplate overnight to ensure full crosslinking. 16 wells were punched into each PDMS disc using a 4 mm disposable biopsy punch (Kai Medical, Inc., USA). The prepared PDMS discs were then autoclaved. Track-etched polycarbonate filters (25 mm diameter, 1.2 μm pores, RTTP02500, Millipore, Watford, UK) were attached in sterile conditions to the bottom of the PDMS disc, covering the 16 punched wells for the culture of the organoids. The polycarbonate membrane and PDMS are both hydrophobic, sealing reversibly yet sufficiently through van der Waals forces. Immediately prior to use, the PDMS organoid culture discs were placed on PDMS separators to create a 2 mm void underneath the polycarbonate membrane in a 6 well plate (Corning) and the respective well was filled with 3 ml MEM (+10% FCS, without phenol red, Fisher Scientific UK Ltd. 11504506). The individual organoid wells were also filled with medium.

### Isolation of embryonic kidneys

For all experiments, kidneys were isolated from E13.5 embryos after humane sacrifice of mice. For the initial experiments, wild-type pregnant CD1 mice (Charles River, Margate, UK) were used. Alternatively, Wt1^+/GFP^ (Wt1^tm1Nhsn^, following Wt1-GFP) (34) male mice were crossed to wild-type CD1 female mice. Experimental animal protocols were performed in accordance with the approved guidelines under a licence granted under the Animals (Scientific Procedures) Act 1986 and approved by the University of Liverpool Animal Ethics Committee. Intact rudiments were either fixed immediately after isolation or cultured on a polycarbonate membrane (RTTP02500) placed on a metal grid and grown in DMEM (+10% FCS, D5796).

### Re-aggregation assay and organoid culture

The embryonic rudiments were processed using an established protocol (5). Briefly, they were pooled, washed 3 times with PBS and dissociated in 3 ml of 1× trypsin in PBS at 37°C for five minutes. Every two minutes, the fragments of the rudiments were gently pipetted up and down to assure complete cellular dissociation of the tissue. The fragments/cells were stabilised with 10 ml DMEM + 10% FCS at 37°C for five minutes and centrifuged to obtain a pellet. The resuspended pellet was then counted by Trypan blue exclusion using a TC20 automated cell counter (Bio-Rad Laboratories, Inc., USA), presenting with an average cell viability of 77% (wild-type) and 69% (Wt1-GFP). For each assay, 1×10^5^ cells were dispensed in 500 μl microfuge tubes and centrifuged at 3000 rpm for two minutes. Each pellet was carefully detached from the tube wall and placed into a PDMS organoid well where the organoids compacted overnight. The organoids were cultured in this setup in a humidified incubator at 37°C and 5% CO_2_ for up to six days (see Table 1 for details on treatment times).

**Table 1:** Time regimes of mouse embryonic kidney organoid and organ treatments.

| Time<br>(d) | Endpoint fixing and staining |  | Functional assay |  | Live imaging |  |
| --- | --- | --- | --- | --- | --- | --- |
|  | Organoid | Kidney | Organoid | Kidney | PNA stain pilot | Wt1-GFP |
| 0 | Re-aggregation | Fix | Re-aggregation | Culture | Re-aggregation | Re-aggregation |
| 1 | Culture |  | Culture | Culture | Culture | Live imaging |
| 2 | Culture |  | Culture | Culture | Culture |  |
| 3 | Culture |  | Culture | Culture | Culture |  |
| 4 | Culture |  | Culture | Functional assay | Live imaging |  |
| 5 | Culture |  | Culture |  |  |  |
| 6 | Fix |  | Functional assay |  |  |  |

### Gel embedded culture of organoids

Spheroids for live imaging were pre-stained in medium containing 30μg/ml rhodamine-labelled Peanut agglutinin (PNA, Vector Laboratories, USA) for 30 mins. Live imaging of organoids was performed on samples that were embedded in a hydrogel cylinder consisting of an agarose-gelatine mix inside a glass capillary. The hydrogels were prepared in sterile conditions. Prior to use, agarose aliquots consisting of low melting agarose (3% in PBS, Sigma A9414) were melted in a heat block at 75°C before cooling down to 38°C. Gelatine aliquots (6% in PBS, Sigma G1890) were heated to 38°C and added to the agarose before embedding the organoids. The appropriate capillary sizes (size 1 ~0.68 mm, size 2 ~1 mm, size 3 ~1.5 mm, size 4 ~2.15 mm) were chosen for the sample to occupy between 1/3 and 2/3 of the agarose diameter. For culturing and imaging, the hydrogel cylinder was partially extruded so that the organoids were located outside the capillary. The imaging chamber was filled with ~12ml of phenol red-free MEM supplemented with 10% FCB, 1% penicillin/streptomycin and 3μg/ml PNA-rh.

### Fixing

Medium was removed from the incubated samples and intact kidneys, pellets and organoids were washed with PBS followed by fixation with 4% paraformaldehyde (PFA) for 30 mins at room temperature and three further PBS washes.

### Clearing

The clearing protocol chosen for this study is based on the CLARITY method (35-38), involving the embedding of the tissue in an acrylamide-based hydrogel followed by the removal of the main scattering components, i.e. lipids. The removal of the lipids also enables a deeper penetration of immunostaining agents, ultimately resulting in optically cleared tissue whilst maintaining structural information.

Hydrogel Preparation: The embedding hydrogel was prepared according to an established protocol (36). The materials used to make up 40 ml of the final monomer solution are the following: 4.7 ml of Acrylamide/Bis (30%, 37.5:1), 100.5mg of VA-044 Initiator (2,2’-Azobis[2-(2-imidazolin-2-yl)propane]dihydrochloride, Alphalabs 017-19362, final concentration 0.25%), 4 ml of 10× PBS (final concentration 1×) and 10 ml of 16% PFA (final concentration 4%) in 21 ml ddH_2_O. The individual components were kept on ice during mixing to avoid polymerisation. The final solution was stored at -20°C until needed.

Hydrogel sample embedding:_Hydrogel monomer aliquots were thawed at 4°C or on ice and then gently mixed to disperse any precipitate. The sample was introduced into the hydrogel monomer solution and kept on ice. They were incubated on a rocker overnight at 4°C for hydrogel infusion. Prior to polymerisation, a layer of peanut oil was added on top of the aliquots and polymerisation was initiated and completed by keeping the samples at 37°C for 3 hours in a heating block. Following polymerisation, the peanut oil was discarded and excess hydrogel was removed from the samples using dissection tools, followed by washing 3× with PBS and storage at 4°C until needed.

Lipid Removal:_The final clearing solution was prepared by dissolving 3.2 g of SDS in 40 ml of ddH2O under agitation and stored at room temperature. The samples were cleared through passive clearing (38) by immersing the samples in the clearing solution and incubating them at 37°C for 48 hours. Finally, the samples were washed 3× with PBST (0.1% Tween in PBS) for 10 mins to remove remaining SDS micelles. The cleared samples were stored in PBST at 4°C until needed.

### Immunofluorescence of fixed organoids

Samples, some cleared and some un-cleared, were blocked for 2 hours at room temperature in blocking buffer (10% goat serum with 0.1% TritonX in PBS). The samples were incubated with primary antibodies diluted in blocking buffer at 4°C using conditions shown in Table 2. Secondary antibodies were diluted at a concentration of 1:1000 in blocking buffer and incubated for 2 hours at room temperature prior to staining with 30 μg/ml PNA for 30 mins. The secondary antibodies used were: Alexa Fluor™ 405 Goat Anti-Rabbit IgG (H+L), Alexa Fluor™ 633 Goat Anti-Mouse IgG_1_, Alexa Fluor™ 488 Goat Anti-mouse (Invitrogen Fisher Scientific).

**Table 2:** List of primary antibodies and stains used.

| Renal structure | Antibody | Supplier | Product | Dilution | Incubation time |
| --- | --- | --- | --- | --- | --- |
| Basement membrane | **Laminin** | Sigma-Aldrich | L9393 | 1:1000 | Overnight |
| Tubular lumen | **Megalin** | Acris | DM3613P | 1:200 | Overnight |
| Ureteric bud incl. tips | **Cytokeratin** | Abcam | Ab115959 | 1:100 | Overnight |
| Metanephric mesenchyme | **Pax2** | Abcam | Ab37129 | 1:200 | Overnight |
| Metanephric mesenchyme | **Six2** | Proteintech | 11562-1-AP | 1:200 | Overnight |
| Podocytes in glomeruli, Metanephric mesenchyme | **Wt1** | Millipore | 05-753 | 1:200 | Overnight |
| Renal podocytes | **Synaptopodin** | Progen | 65194 | 1:1 | Overnight |
| Renal podocytes | **Nephrin** | Abcam | Ab72908 | 1:180 | Overnight |
| Basement membrane | **Rhodamine labelled Peanut Agglutinin (PNA)** | Vector Laboratories | RL-1072 | 20-30 µg/ml | 30 mins |

### Functional (organic anion transport) assay

All assays were performed on live cultures. The intact kidney rudiments were cultured at the air liquid interface for 4 days following dissection. The organoids were cultured under medium immersion for 6 days following dissection. The samples cultured at the air liquid interface were gently removed from the membranes before starting the assay.

The samples were incubated for 1 hour at 25°C with PBS containing 1μM 5(6)- carboxyfluorescein (6-CF) and 20 μg/ml of PNA (Vector Laboratories). For the control samples, 2 mM probenecid was also added to the solution to inhibit the organic anion transport. After incubation, the samples were washed with ice cold PBS for 10 mins followed by an incubation with 8mM probenecid in PBS for 15 mins to arrest any transport. The functional assay was concluded by two PBS washes and samples were then embedded in 1. 5% agarose for imaging.

### Imaging

Fixed and stained samples were embedded in 1.5% agarose columns using appropriately sized glass capillaries. The glass capillaries were inserted into the imaging chamber of the microscope and the agarose column was extruded into the chamber filled with PBS. The light sheet fluorescence microscope used was a Zeiss Lightsheet Z.1 microscope, fitted with 10× illumination objectives and an achromatic 20× detection objective. The laser wavelengths used were 405 nm, 488 nm, 561 nm, 638 nm and the laser intensity was kept to a minimum. Green fluorescent beads, 30 nm small beads (polystyrene latex beads (470/505), Sigma-Aldrich Corp., USA) or 1 μm large beads (Fluospheres (505/515), Thermo Fisher Scientific Inc., USA), were used as fiduciary markers.

### Data Analysis

Data analysis was performed using the Zen analysis software (Carl Zeiss AG, Germany) as well as Fiji (ImageJ) (39). Cell tracking was performed using the TrackMate plugin (v.2.8.2) for Fiji (40) and three-dimensional visualisations of the data were generated using the 3D Viewer (v.) plugin for Fiji (41). The TrackMate (42) plugin for Fiji is a single particle tracking tool that identifies spots, i.e. cells, in every frame and the trajectories of cells are reconstructed by assigning an identity over these frames in the shape of a track. The positional data and numerical features generated using TrackMate were further processed and displayed using Matlab (R2014a, The Mathworks Inc., USA). Violin plots were generated using a script obtained via https://github.com/bastibe/Violinplot-Matlab. The Mann-Whitney U procedure for statistical testing was run in Matlab. Multiview reconstruction was performed using the MultiView Reconstruction plugin in Fiji (43, 44). Object segmentation and volume calculation of the segments was done in Imaris (8.4.2, Bitplane AG, Switzerland).

### Data Accessibility

The raw imaging data that this manuscript is based on has been deposited in the “Image Data Resource” and is publicly available as dataset idr0038 under CC-BY 4.0 licence, providing full access. The data DOI is 10.17867/10000110.

## Results and Discussion

We have adapted the kidney rudiment re-aggregation assay by creating embryonic renal organoids suitable for light sheet fluorescence microscopy (Figure 1). These organoids and also intact kidney rudiments were imaged with a light sheet fluorescence microscope after end point fixing and staining to determine the organotypical development of structures in the organoids. Furthermore, we aimed to image the tissues *in vitro* over the long-term to track labelled cells during the culture period, and subsequently analyse the tracking data.

**Figure 1.**
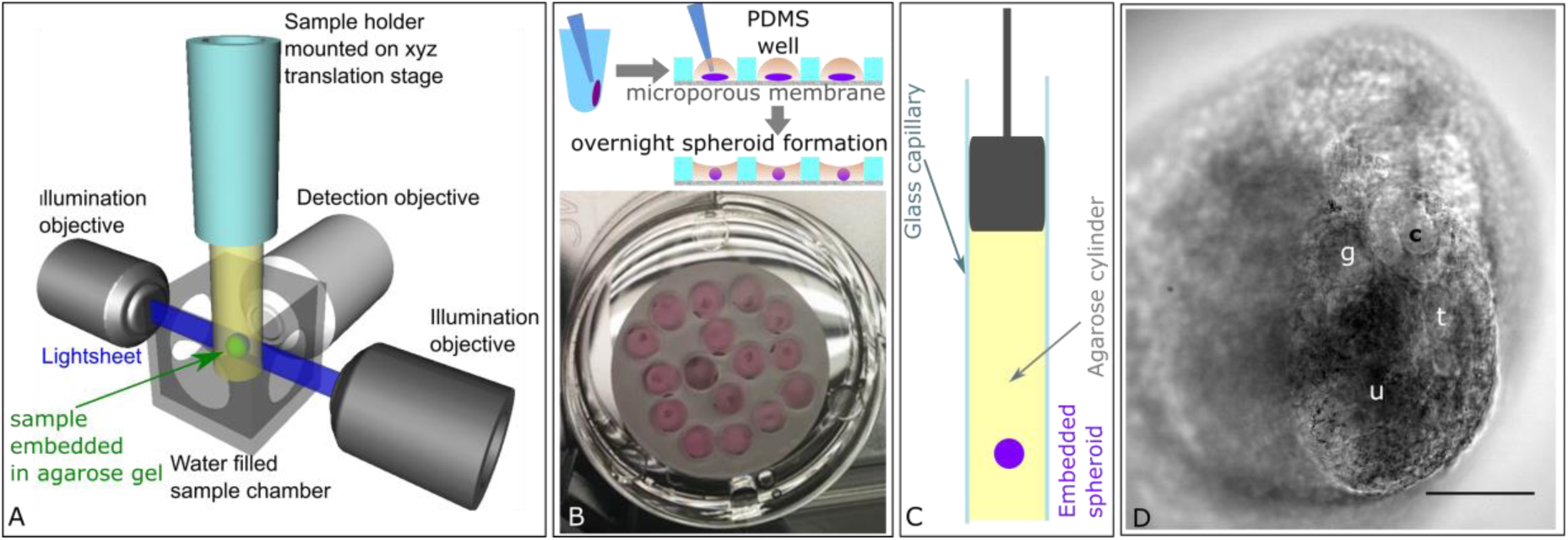
Experimental setup. *A) Light sheet fluorescence microscopy*. A single plane of the specimen, embedded in an agarose gel cylinder, is illuminated by a light sheet with the detection located orthogonally to the illumination. Only the plane that is observed is illuminated, resulting in vastly reduced photo bleaching and phototoxic effects. The specimen can be moved and rotated in the fluid filled chamber and through the stationary light sheet. *B) Embryonic kidney organoid assay.* To generate organoids, the renal pellets were removed from the centrifuge tubes and placed in organoid wells in the PDMS disc. Overnight, the pellets compacted into organoids. C) For light sheet imaging, the organoids were embedded into an agarose cylinder inside a glass capillary. D) Transmitted light image of a renal tissue spheroid embedded in agarose. The focal plane shows several renal structures including a tubule (t), ureteric bud tip (u), comma shaped body (c) and glomerulus (g). Scale bar = 100 μm.

### Optimising protocols for generation of re-aggregated embryonic renal organoids for light sheet microscopy

For post-fixation imaging of the developing kidney structures, we generated re-aggregated organoids from embryonic kidneys at E13.5 following previously published protocols (5, 6, 45, 46). The pellet was transferred into an organoid well fabricated in a PDMS disc, where the size of the well and its surface hydrophobicity promoted pellet compaction overnight, leading to the formation of an organoid that did not attach to the well (Figure 1B). The micro-porous membrane at the bottom ensured the exposure of the organoid to growth factors in the medium, thus supporting its growth and enabling the recapitulation of developing structures (Figure 1D). For live imaging in a light sheet fluorescence microscope, we embedded the samples in an agarose-gelatine hydrogel using glass capillaries and cultured them for up to 15 hours while recording (Figure 1C, D). The agarose-gelatine hydrogel mixture supported the development of nephron structures and the addition of a fluorescent vital stain. Turning the gel column around the gravitational axis, we could repeatedly image the organoids from various angles.

Throughout the 6 day culture period, we observed a reduction in size of the organoids, both in those cultured in medium and in those cultured in the agarose-gelatine hydrogel (Figure 1D and SI Figure 1). On day 4, the organoids cultured in an agarose-gelatine gel shrank to ~57% of the size recorded 24 hours after re-aggregation, i.e. after organoid formation (n=3).

### Renal marker analysis in fixed embryonic kidneys and re-aggregated renal organoids

To demonstrate that the culture method above resulted in 3D kidney organoids with appropriate organotypic structures, we probed PDMS-cultured organoids with a panel of markers (Table 1). As a comparison and control of the specificity of the markers, we included intact embryonic kidney rudiments (E13.5). Maximum intensity projections of stained embryonic kidneys and organoids allowed a quick comparison of their three-dimensional arrangement since maximum intensity projections plot the signal of the respective markers distributed throughout the samples onto one plane.

Light sheet fluorescence microscopy imaging of laminin-stained embryonic kidneys and organoids showed an arrangement of basement membranes which could indicate both the development of the ureteric bud and/or tubules (SI Figure 2). The proximal tubule-specific receptor megalin was rarely detectable in the embryonic kidneys at E13.5 (47) (SI Figure 2B),i. e. laminin^+^ structures probably indicate ureteric bud and immature proximal tubules. In the more mature organoids, megalin was expressed within the lumen of the majority of laminin+ structures (SI Figure 2C and D), consistent with the development of epithelial tubules (5). Cytokeratin-stained embryonic kidneys revealed the presence of collecting duct trees with several branches ending in ureteric buds (SI Figure 3A and E) (48). Similarly, branched though less organised cytokeratin+ structures could be identified in 6-day old organoids (SI Figure 3C and G).

We included the ureteric bud in the generation of the single cell suspension. As a result, the ureteric bud cells formed independent epithelial foci which did not lead to the formation of a sole ureteric tree. To circumvent this problem, alternative re-aggregation protocols start from the separation of the metanephric mesenchyme from the ureteric bud, where the ureteric bud is either discarded or cultured separately and then re-added to the metanephric mesenchyme mix at the beginning of the culture period (2, 49). This method results in a more physiological renal tissue aggregated around one ureteric bud tree.

For the detection of cap mesenchyme we used Pax2 antibody staining, which was arranged around basement membrane-bound ureteric bud structures in intact kidneys and reaggregated organoids (SI Figure 4). Pax2 was highly expressed in the condensing mesenchyme but was also detectable within the cells of the tubules (SI Figure 4E and G) (50).In both the intact embryonic kidneys and organoids, the cap mesenchyme consisted of 4-5 cell layers (SI Figure 4E-H, SI Video 1). Re-aggregated organoids also expressed the nephron progenitor marker Six2 in the cap mesenchyme (SI Figure 5).

Light sheet fluorescence microscopy has the advantage to image much deeper into the tissue than traditional confocal microscopy. However, the penetration of antibodies was limited and this was particularly the case for the laminin antibody. To exploit the deep imaging capability of the light sheet microscope, the pan-epithelial lectin stain PNA was used as an alternative marker for basement membranes. The stain only required one rapid staining step, penetrated fully through the samples, and was therefore used throughout this study to highlight the networks of ureteric bud branches and tubules within embryonic kidneys and organoids (SI Figure 3 and SI Figure 4). SI Figure 5B shows a panel of single focal planes of a stained organoid at increasing depth. Only the endogenous Wt1-GFP and the PNA signals were detected throughout the spheroid, while the antibody signals (Six2, Synaptopodin) were limited to 75 μm from the organoid surface.

### Detecting glomerular maturation in spheroidal re-aggregated kidney organoids

To track developmental cell changes in renal structures over time in the light sheet microscope, we used embryonic kidneys from Wt1-GFP knock-in reporter mice where GFP is expressed under control of the endogenous Wt1 promoter. Wt1 is dynamically expressed throughout kidney development in the metanephric mesenchyme, cap mesenchyme, renal vesicles, S- and comma-shaped bodies and podocytes (51, 52). We found that Wt1 was expressed in the cap mesenchyme, renal vesicles, S and comma-shaped bodies in both intact, fixed E13.5 kidneys and organoids cultured in the PDMS-polycarbonate membrane well for 6 days (SI Figure 6). Due to the difference in the developmental stages of the embryonic kidneys and organoids, we only observed a strong signal in the maturing podocytes of the glomeruli in the organoids. In maximum intensity projection and single focal plane images of two organoids, fixed and stained after 6 days of culture, several bright Wt1+ structures were visible that co-labelled with the podocyte markers nephrin or synaptopodin, respectively (Figure 2A and H).

**Figure 2:**
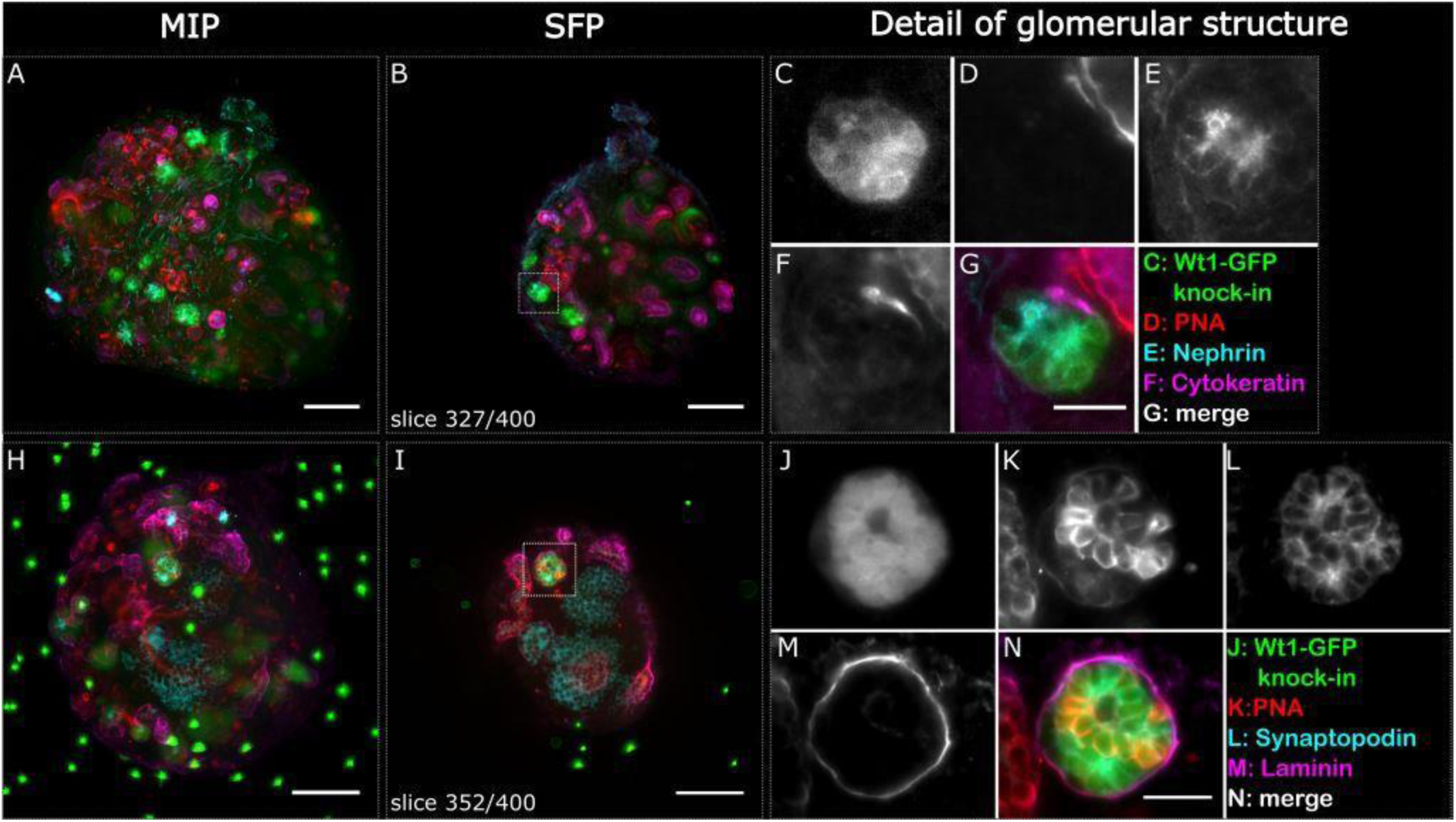
Glomerular structures stained positively for podocyte markers indicating phenotypic maturity in re-aggregated embryonic renal organoids after 6 days of culture. Two whole organoids (A-G and H-N) were imaged in the light sheet microscope and are depicted as maximum intensity projections (A, H) and respective selected single focal planes (B, I). Glomerular structures are highlighted with a dashed square in the single focal plane, and detailed images are shown in C-G and J-N respectively. The Wt1-GFP structure in C also stained positive for nephrin (E), and the Wt1-GFP structure in J stained positive for synaptopodin (L). Cytokeratin highlights the various ureteric bud foci, and PNA and laminin the basement membranes of the ureteric bud and nephric tubules. The sample shown in H-N contains fluorescent beads as reference points for multi view reconstruction of the dataset. The organoids were not cleared. Scale bars in A, B, H and I: 100 μm, G: 10 μm, N: 25μm.

Nephrin (Figure 2E) is a slit diaphragm protein and part of the renal filtration barrier essential for maintaining normal glomerular permeability. In kidney organoids cultured for 6 days, the marker is present on the apical surface of Wt1-expressing podocytes (Figure 2C). In another staining panel, the Wt1+ clusters (Figure 2J) co-labelled with synaptopodin (Figure 2L). The expression of synaptopodin in podocyte foot processes is differentiation-dependent, indicating advanced cytoskeleton development and is considered an important marker of the phenotypic maturity of podocytes. These results indicate normal organotypic development of glomeruli in the re-aggregated kidney organoids (53).

### Physiological tubular transport in re-aggregated kidney organoids

To test the physiological function of the developed tubules in 6-day old spheroidal reaggregated kidney organoids, we employed an organic anion transporter assay. Specifically, we assessed whether the tubule cells were capable of organic anion transporter-dependent uptake of the organic anion mimic 6-Carboxyfluorescein (6-CF). For comparison, we included intact embryonic kidney rudiments collected at E13.5 that had been cultured for 4 days at the air-liquid interface. Since organoids lag behind in their developmental progress due to the reaggregation process, the intact rudiments were cultured for 4 days to reach a maturity similar to 6-day old organoids. Our analysis showed that in both the cultured embryonic kidneys and the re-aggregated organoids, 6-CF was transported into the cells (Figure 3A and C). Furthermore, the organic anion transporter-based 6-CF uptake could be successfully prevented using the inhibitor probenecid (Figure 3B and D), both in organoids and cultured embryonic kidneys. These findings demonstrate active organic anion transport in renal tubules formed in the spheroid re-aggregated embryonic kidney organoids. Our results are in agreement with previous organic anion transporter assays that have shown that tubules, developed in re-aggregated engineered kidney tissue, exhibit transport function, thus mimicking physiological processes (6, 54).

**Figure 3:**
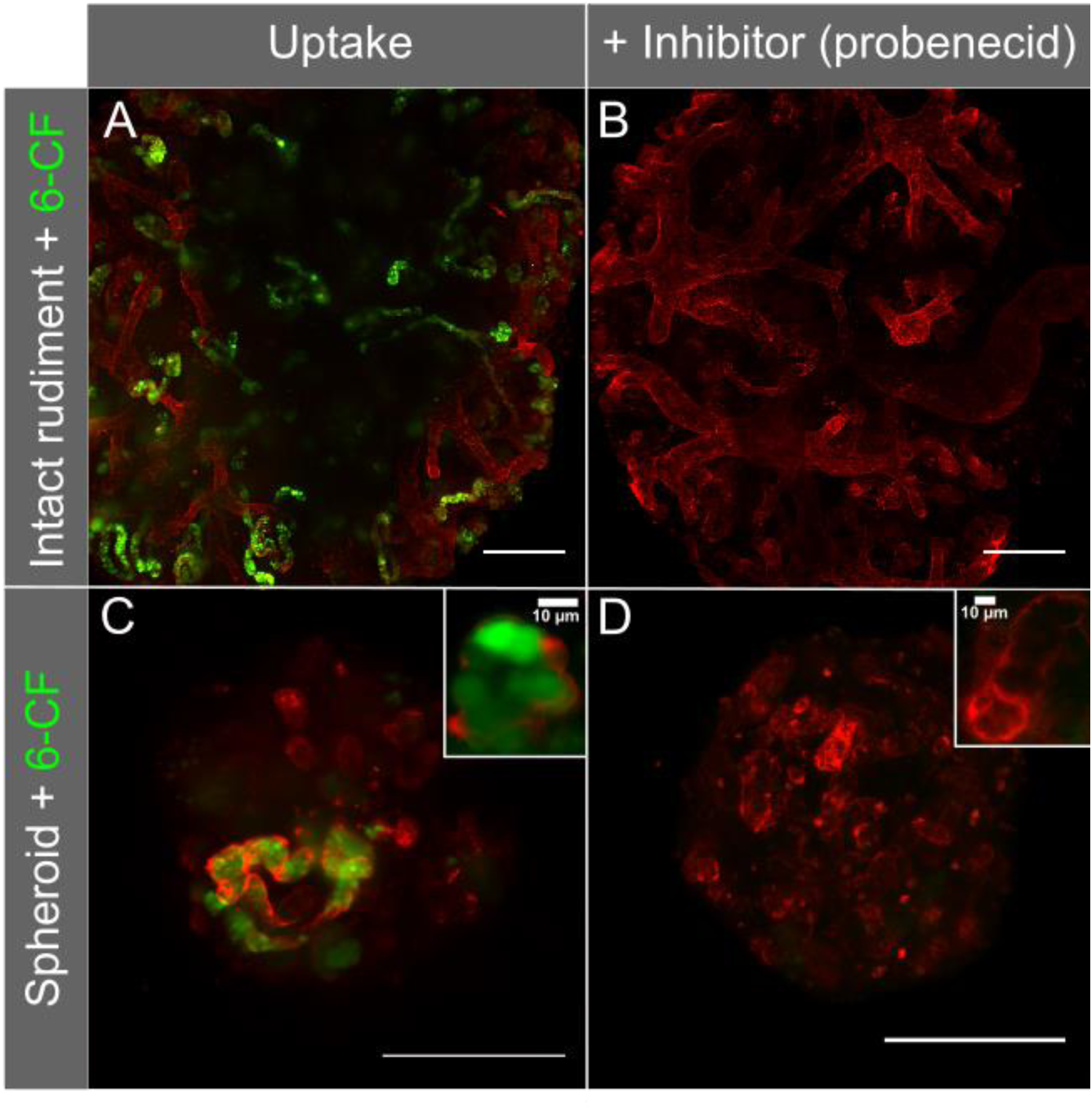
Specific uptake of the fluorescent anion 6-Carboxyfluorescein with and without the organic anion transport inhibitor probenecid. The basement membranes of the tubules in the cultures were highlighted with rhodamine-conjugated PNA (red). Uptake was investigated in E13.5 mouse rudiments cultured for 4 days on a micro-porous membrane at air-liquid interface, and re-aggregated organoids cultured for 6 days immersed in a PDMS well system with a bottom layer of micro-porous membrane. The images are maximum intensity projections of z-stacks. Scale bar = 200μm.

### Live imaging and cell tracking in spheroidal organoids cultured inside hydrogel

We could demonstrate that organotypic structures can form in embryonic kidney organoids after re-aggregation. Aiming to record cell migration and differentiation we imaged the organoids live, by culturing them inside hydrogel cylinders surrounded by medium within the light sheet fluorescence microscope. In a pilot experiment, we generated wild-type embryonic mouse kidney organoids followed by 4 day culture in the PDMS wells, before embedding in the hydrogel cylinders. We chose a 4-day old organoid because based on our previous results we expected that ureteric bud foci and nephric tubules had been established and their basement membranes should stain readily, providing structural context for the spheroid. The sample chamber was filled with medium containing a low concentration of PNA, for continuous staining during live cell imaging. The compaction of the organoid over time was clearly visible (SI Figure 7A). The dye gradually diffused into the tissue. Initially, the PNA signal seemed indiscriminate and no structures were recognisable up until 4 hours into time series (SI Video 2). Subsequently, live samples were pre-stained prior to imaging to avoid the delay in structure detection. Throughout the time series individual, labelled cells could be identified and tracked, including cell integration into a tubule (SI Figure 7B, SI Video 2). The development of renal structures in 4-day old organoids can therefore be visualised over a 13 hour time period in light sheet live cell imaging.

To investigate the cell dynamics during nephron formation in live organoids, we cultured re-aggregated renal organoids from Wt1-GFP reporter mice in PDMS dishes for 24 hours (Table 1), pre-stained with PNA, and then embedded within the hydrogel cylinders for continued culture and live imaging. During the imaging period and while structures developed, the Wt1-GFP cells continually expressed GFP, allowing the automated tracking of cells over time (SI Video 3 and 4). Typically, PNA-labelled structures were visible at the start of the imaging series, i.e. 24 hrs following cell re-aggregation (Figure 4A for “Experiment 1”, T = 0.5 hr and Figure 5A, T = 0 hr, and SI Video 5 for “Experiment 2”). For both experiments, Wt1-GFP cells were grouped around PNA-labelled basement membrane bound structures whereas the inner parts of the structures were entirely devoid of Wt1-GFP cells throughout the imaging periods of 15 hours. This pattern reflects the situation *in vivo*, where the ureteric bud in mouse embryonic kidneys is surrounded by Wt1^+^ cap mesenchyme cells. Most of the cells surrounding the ureteric bud in Figure 4A expressed GFP but some did not (arrows). As the cells are not labelled with any other marker, it is unclear whether they started expressing GFP at later time points or moved away from the cap mesenchyme. Figure 4B shows a threedimensional render of another area within the same organoid, containing a second basement membrane bound structure void of GFP expressing cells but surrounded by them over a period of eight hours.

**Figure 4:**
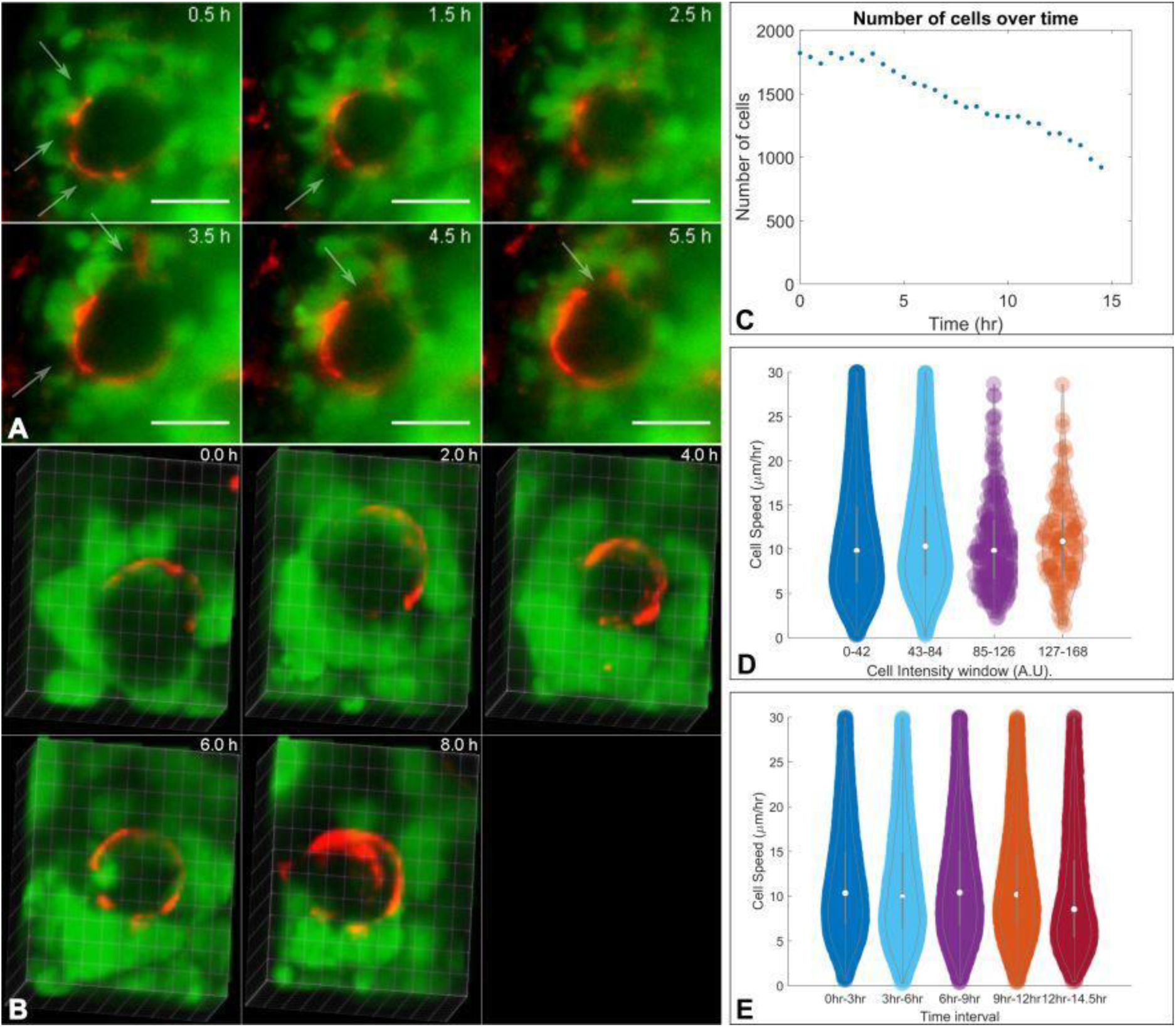
Live imaging of renal organoids using light sheet fluorescence microscopy A) Detailed views of representative single focal plane images showing Wt1-GFP cells (green) grouped around a ureteric bud structure labelled with PNA (red). Over the depicted period of 5 hours, Wt1-GFP cells were entirely absent from the ureteric bud. Most of the cells surrounding the ureteric bud structure expressed GFP but there were also some dark cells (arrows). Scale bar 25 μm. B) shows a three-dimensional render of another basement membrane surrounded structure void of Wt1-GFP cells within the same organoid over a time period of 8 hours. C-E show a selection of parameters analysed following automated cell tracking. (C) The overall number of GFP expressing cells declined slowly over time. (D) Cells were grouped into different intensity windows to compare the speed at which they moved, revealing no notable difference. The cell intensity distribution is skewed towards low intensities. (E) The cell speed does not change much over the 15 hour imaging period.

As we captured the three-dimensional arrangement of living GFP-expressing cells within the organoids every 30 minutes for 15 hours with the light sheet microscope, we could track their positions over time. The single particle tracking tool TrackMate identifies cells and their tracks throughout the time series, generating a range of data sets. We plotted the cell distribution in three dimensional space and calculated the median distance of each cell to the centre of mass of the organoid. This enabled us to estimate the size of the organoid and its change over time, thus quantifying compaction (SI Video 6). Some of the quantitative data derived from Experiment 1 is represented in Figure 4C-E. The number of Wt1-GFP cells declined over time by 50% at 14 hrs (Figure 4C), giving some indication that de-differentiation and/or cell death occur. The median cell intensity also declined during the time series (data not shown) and viewing the time series in detail shows that there is cell death of Wt1-GFP expressing cells. Quantitative parameters like the cell speed and median intensity can be correlated at the single cell level (Figure 4D). Parameters like the track median speed can be plotted as a histogram over the whole imaging period but more information can be derived from windowing the speed of cells (Figure 4E). The violin plot shows that the cell speed distribution remains constant throughout the first 12 hours followed by a slight drop during the 12-14.5 hr interval.

To evaluate reproducibility we repeated the time series experiment, setting Experiment 2 up identically to Experiment 1. Detailed views of a single focal plane of Experiment 2 over 23.5 hours are shown in Figure 5A. As in Experiment 1, a basement membrane-bound structure surrounded by Wt1-GFP expressing cells is clearly identifiable. Figure 5B shows a 3D segmentation version of the same structure (arrow). The volume was calculated from the segmentation and increased steadily (Figure 5C). The shape of the ureteric bud changed over time, ultimately resembling two tips emanating from the original spherical shape. The mean cell diameter remained constant over time (Figure 5D) but the number of cells decreased like in Experiment 1.

**Figure 5:**
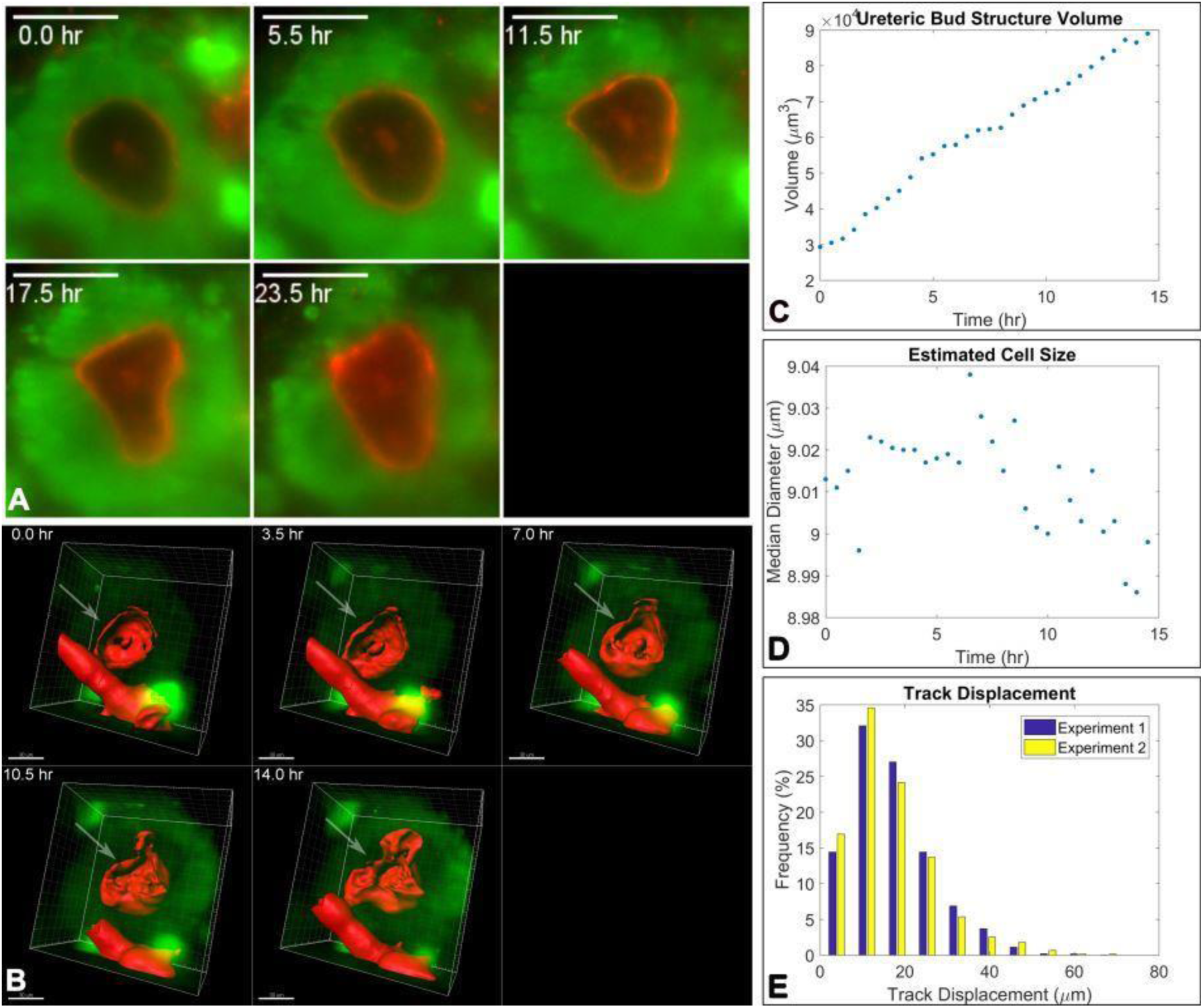
Live imaging of renal organoids using light sheet fluorescence microscopy. A) Detailed views of representative single focal plane images showing Wt1 -GFP cells (green) grouped around a ureteric bud structure labelled with PNA (red). Over the depicted period of 23.5 hours, Wt1-GFP cells were entirely absent from the ureteric bud. Every cell surrounding the ureteric bud structure appeared to express GFP. Scale bar 25 μm. B) Three-dimensional segmentation of the ureteric bud structure shown in A). The ureteric bud segment is indicated with an arrow. The rod-shaped segment at the bottom is a piece of dust integrated in the organoid. Scale bar: 30 μm. C) represents the volume increase of the ureteric bud structure throughout the time series. D) shows that the cell diameter does not change over time. E) shows a comparison of the track displacement histograms of the two time series discussed in this and the previous figure, which are similar.

Quantitative tracking data enables quantitative comparison between experiments, which was not possible previously. The mean cell diameter of Wt1-GFP cells determined via TrackMate is similar for both time series (9.7 μm and 9.0 μm). Figure 5E shows the track displacement histograms of both experiments that were set up identically on different days. The distributions have a similar shape with the mean of the track displacement of 15.4 μm (n = 3026) for Experiment 1 and 14.2 μm (n = 1423) for Experiment 2. Therefore, obtaining quantitative data through live imaging in the light sheet fluorescence microscope, of the developmental processes that can be tracked, opens up an avenue to statistical analysis.

## Conclusion

We show that we can reliably create renal tissue organoids from disaggregated mouse embryonic kidneys using a custom construct made of PDMS. We created a customizable multi-well enabling the formation of up to 16 organoids in parallel, hence reducing the associated costs and intra-experimental variability. This set-up also holds potential to create higher-throughput platforms for drug testing and additional studies. The generated renal and chimeric tissue organoids were imaged with light sheet fluorescence microscopy. Structure and tubular function were preserved *in vitro*. Within the organoids imaged cells long-term, and subsequently tracked the cell fate and generated previously unavailable quantitative cell data.

## Acknowledgments

This work was supported by the UK Regenerative Medicine Platform (MR/K026739/1). We acknowledge the Liverpool Centre for Cell Imaging (CCI) for provision of imaging equipment, in particular the Zeiss Lightsheet Z.1 funded by BBSRC Alert13 grant number BB/L014947/1 and technical assistance. I.S. was funded by the Nephrotools FP7 Marie Curie training grant, and Alder Hey Children’s Kidney fund. We would also like to thank Thomas Wilm for help with the genotyping of the mice, and Marco Marcello and David Mason for help with technical assistance and image analysis support.

## Author Contributions

M.H.: concept and design, collection and/or assembly of data, data analysis and interpretation, manuscript writing, final approval of manuscript, I.S.: conception and design, collection and/or assembly of data, data analysis and interpretation, manuscript writing, B.W., P.M., and R.L.: conception and design, financial support, data analysis and interpretation, manuscript writing, final approval of manuscript.

